# Contrasting the development of larval and adult body plans during the evolution of biphasic lifecycles in sea urchins

**DOI:** 10.1101/2024.04.30.591978

**Authors:** Brennan D. McDonald, Abdull J. Massri, Alejandro Berrio Escobar, Maria Byrne, David R. McClay, Gregory A. Wray

## Abstract

Biphasic lifecycles are widespread among animals, but little is known about how the developmental transition between larvae and adults is regulated. Sea urchins are a unique system for studying this phenomenon because of the stark differences between their bilateral larval and pentaradial adult body plans. Here, we use single cell RNA-sequencing to analyze the development of *Heliocidaris erythrogramma* (*He*), a sea urchin species with an accelerated, non-feeding mode of larval development. The sequencing time course extends from early embryogenesis to roughly a day before the onset of metamorphosis in *He* larvae, which is a period that has not been covered by previous datasets. We find that the non-feeding developmental strategy of *He* is associated with several changes in the specification of larval cell types compared to sea urchins with feeding larvae, such as the loss of a larva-specific skeletal cell population. Furthermore, the development of the larval and adult body plans in sea urchins may utilize largely different sets of regulatory genes. These findings lay the groundwork for extending existing developmental gene regulatory networks to cover additional stages of biphasic lifecycles.

**Summary statement:** Analysis of a new single cell transcriptomic atlas of sea urchin development reveals the rapid evolution of larval cell type trajectories and provides a candidate list of regulators for adult rudiment development.

## Introduction

Most metazoan phyla contain species with biphasic lifecycles, also known as indirect development (Formery & Lowe, 2023; Moran, 1994; Nielsen, 2009; Strathmann, 1985; Thorson, 1950). Species that use this life history strategy first develop from embryos into a larval body plan, and then undergo metamorphosis to transition into a separate adult body plan. Little is known about how the transition between the larval and adult body plans is regulated at the molecular level. Larval development of phyla such as the hemichordates and annelids is modeled to be controlled by the bilaterian anterior patterning network, while the delayed activation of the trunk patterning network is proposed to initiate adult development (Formery & Lowe, 2023; Gonzalez et al., 2017; Lacalli, 2005; Martin-Zamora et al., 2023).

This model does not fit with sea urchin development. These animals have highly derived body plans due to the pentaradial symmetry of their adults (Smith, 2008). They evolved from a bilaterally symmetrical ancestor, and their larvae retain this symmetry form (Peterson et al., 2000; Sumrall & Wray, 2007). Thus, sea urchins undergo a developmental transition between larvae with bilateral symmetry and juveniles with pentaradial symmetry when they metamorphose. A recent study found that the adults of sea stars, a close relative of sea urchins, do not employ the bilaterian trunk regulatory network to pattern the ectoderm (Formery et al., 2023), unlike hemichordates and annelids (Gonzalez et al., 2017; Martin-Zamora et al., 2023). This suggests that echinoderms may use mechanisms different from other bilaterians to control the transition between the larval and adult body plans. To obtain a better understanding of the relationship between larval and adult pattern regulation, a needed starting point is to see whether the well-characterized gene regulatory network (GRN) that controls embryonic and larval development in sea urchins (Davidson et al., 2002; Peter & Davidson, 2011; Rafiq et al., 2012; Sethi et al., 2009; Su et al., 2009) also plays a role in patterning the adult.

Late larval development and adult morphogenesis in most sea urchin species is difficult to study because development of the adult rudiment takes place over several weeks during the larval phase. This handicap is much reduced with selection of a model species that rapidly transits from the larval to adult body plan. The Australian sea urchin *Heliocidaris erythrogramma* (*He*) employs a lecithotrophic developmental strategy and enters metamorphosis about four days post fertilization (Raff, 1992; Williams & Anderson, 1975). This life history strategy is enabled not by feeding on plankton, but rather by providing sufficient energy content in its eggs to support the juvenile until it can feed on its own. In contrast, sea urchins with planktotrophic larvae (planktotrophs) require feeding on phytoplankton for an extended time to acquire the energy supply necessary to build the rudiment before metamorphosis (McEdward & Janies, 1997; Strathmann, 1985). Although sometimes termed a “direct” developer, *He* has retained many characteristics of the sea urchin biphasic lifecycle and constructs a partial, bilaterally symmetrical larval body (Emlet, 1995). Thus, *He* can be used as a model for studying the transition between the larval and adult bodies within a manageable experimental timeframe.

*He* is also a useful model for understanding how biphasic lifecycles can evolve over short periods of time. Evolutionary transitions between different modes of larval development are common across marine invertebrate taxa, despite the extensive changes that often need to occur to embryonic development and larval morphology (Strathmann, 1978, 1985; Wray, 1996; Wray & Raff, 1991). Identifying the regulatory mechanisms that control the shift between developmental modes has been difficult, since species with different modes are often distantly related. The distinct advantage of studying *He* is that it is closely related to *H. tuberculata* (*Ht*), a sea urchin with the planktotrophic mode of development (Wray, 2022). Phylogenetic evidence suggests that *He* is separated by only about five million years of evolutionary divergence from *Ht* (Hart et al., 2011; Smith et al., 1990; Zigler et al., 2003). This, along with strong evidence that planktotrophic development is ancestral for sea urchins (McEdward & Miner, 2001; Wray, 1996), implies that *He*’s lecithotrophic developmental process is a derived evolutionary feature (Raff, 1992; Smith et al., 1990). At the same time, *He* and *Ht* adults have highly similar morphologies (Byrne & Ohara, 2017), making them a good model for determining how different larval developmental processes can lead to the same adult body plan.

Here, we use *He* as a model for studying the evolution and development of biphasic lifecycles from two perspectives. First, we advance an understanding on how larval development changed during the evolution of lecithotrophy in *He*. Second, we focus on the regulation of larval and adult body plan development in *He* and compare this to the same processes in species with planktotrophic development. To address these objectives, we performed single cell RNA sequencing (scRNA-seq) for *He* embryos and larvae at 12 time points from 6 to 60 hours post fertilization (hpf). The final atlas combines scRNA-seq data from five new time points of larval stages, along with seven embryonic time points that are the focus of a complementary paper (Massri et al., 2024). Temporal scRNA-seq atlases have been generated for several sea urchin species with planktotrophic development (Foster et al., 2020; Massri et al., 2021; Paganos et al., 2021), but these atlases stopped well before metamorphosis. By contrast, the final time point in the *He* atlas is close to the onset of metamorphosis, which is approximately 96 hpf in *He* larvae raised at 23°C. At 60 hpf, *He* larvae possess a highly developed adult rudiment, including partially formed primary podia and the early water vascular system (Fig. 1) (Minsuk & Raff, 2002). We describe several cell types in *He* with unique transcriptional trajectories compared to those in species with planktotrophic development. In particular, we focus on the loss of a larva-specific skeletogenic cell population, the emergence of an undifferentiated cell population, and the accelerated diversification of neural subpopulations. Furthermore, we compare gene expression patterns between early and late stages of the time course and find that genes from the sea urchin embryonic GRN may play a reduced role in controlling adult morphogenesis. We then leverage the presence of cells from adult tissues in our dataset to generate a list of candidate transcription factors that may play key roles in patterning the adult rudiment. Altogether, the extensive temporal coverage of our dataset makes it a valuable resource for addressing questions about larval and adult morphogenesis in sea urchins and perhaps other species with biphasic lifecycles.

**Figure 1:**
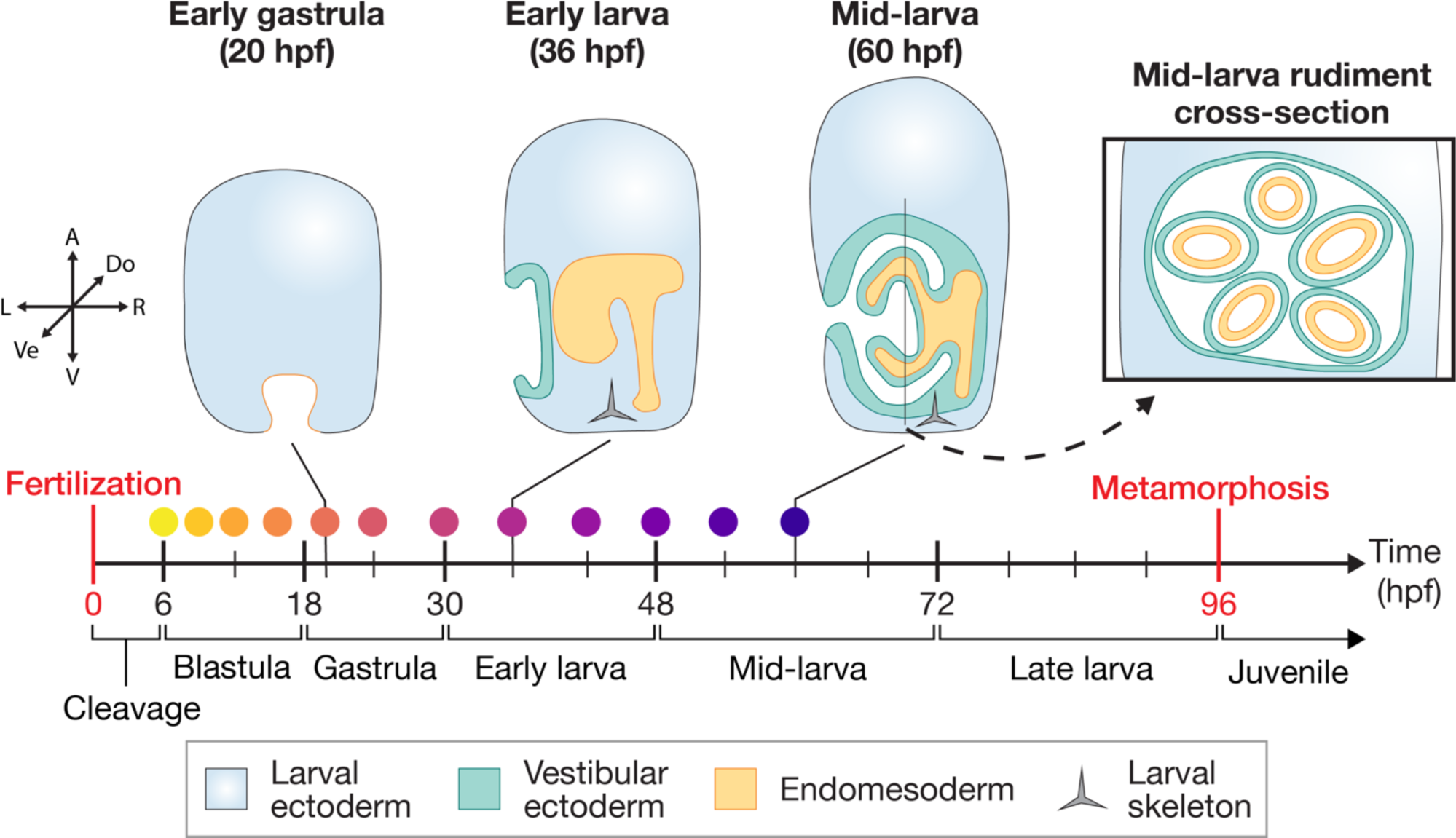
Overview of larval development in *Heliocidaris erythrogramma*. Diagrams of several larval stages are shown at the top, which are vertical cross-sections through the left-right plane of the larva, except for the right-most diagram, which is a cross section of the adult rudiment through the dorsal-ventral plane of the larva. Note that not all tissue types are shown. The timeline below the diagrams highlights the time intervals (in hours post fertilization, hpf) corresponding to key stages in *He* development. The time intervals for each stage are based on raising *He* embryos and larvae at 23°C. Colored dots above the timeline correspond to the scRNA-seq samples collected for this study (see Fig. 2A). The three-dimensional coordinate plane diagram on the left represents the positional axes of the embryo, with labels for the poles of each axis: A, animal; V, vegetal; Do, dorsal; Ve, ventral; L, left; R, right.

## Results

### A temporal scRNA-seq atlas of development in *He* reveals the diversity of cell types in larval sea urchins

Here, we expand upon a temporal analysis of embryonic development in *He* (Massri et al., 2024) by adding scRNA-seq data for five additional time points of larval stages, resulting in insights into formation of the adult rudiment. The final atlas contains 12 time points from 6 to 60 hpf and covers development in *He* from cleavage to the mid-larva (Fig. 1). Analysis of the dataset began by merging cell-level gene expression data from each of the time points into the same object using the scRNA-seq analysis tool Seurat (Hao et al., 2021). After filtering and normalization, there were 49,896 cells across all time points in the dataset. Cells in each time point had an average of 500-1500 genes per cell and 1000-3000 UMIs per cell (Fig. S1). We then used the uniform manifold approximation projection (UMAP) algorithm to project the cells into two-dimensional space for visualization (Fig. 2A) and performed Louvain-based clustering to group cells with similar transcriptional identities. This returned 57 distinct cell clusters, to which we assigned 18 cell type identities (Fig. 2B) based on known marker genes from studies in other sea urchin species (Fig. S2; Table S1). We also performed hybridization chain reaction *in situs* (HCRs) to verify the spatial expression patterns of prominent marker genes from the scRNA-seq atlas. Using this information, we were able to recover most of the cell types that are known to be present in the larvae of sea urchins with planktotrophic development (Massri et al., 2021). For example, we annotated *Pks1* expressing cell clusters as pigment cells (Calestani et al., 2003), which aligns with the HCR expression of *Pks1* in cells scattered throughout the ectoderm in 56 hpf *He* larvae (Fig. 2C). The HCR expression patterns of *Onecut* and *FoxQ2* (Fig. 2D-E) in the ciliary band and animal pole domain of 56 hpf larvae, respectively, also align with known expression patterns in species with planktrophic development (Poustka et al., 2004; Yaguchi et al., 2008).

**Figure 2:**
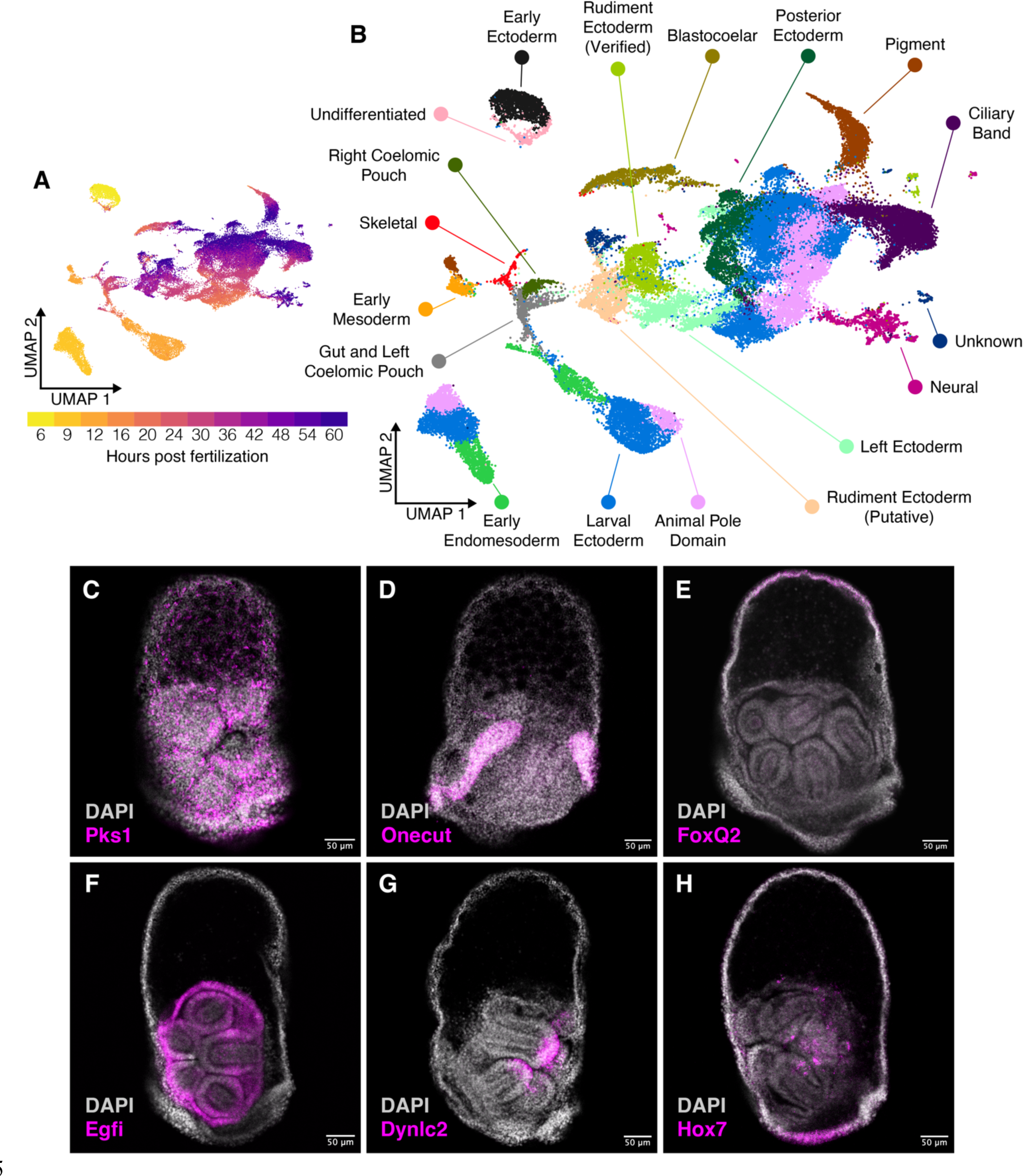
Cell type identification and trajectories in a temporal scRNA-seq atlas of development in *He*. (A) UMAP plot that projects the gene expression profiles of each cell into two-dimensional space, colored by sample time point. (B) UMAP that colors each cell by its cell type identity based on the expression of known cell type marker genes. (C-H), HCR micrographs in 56 hpf *He* larvae for six marker genes that were used to identify cell types in the scRNA-seq atlas. (C) *Pks1* is a marker for pigment cells, (D) *Onecut* is a marker for the ciliary band, (E) *FoxQ2* is a marker for the animal pole domain, (F) *Egfi* is expressed in the rudiment ectoderm, (G) *Dynlc2* is expressed at the tips of the primary podia, and (H) *Hox7* is expressed in the posterior ectoderm and inside the rudiment.

Not all clusters aligned with the profiles identified in previous scRNA-seq studies. Most of these contained cells from later time points, suggesting that the previous scRNA-seq atlases failed to capture these late larval cell types. We used HCRs to map these clusters onto discrete tissues in 56 hpf larvae. For example, one transcript of *Egfi*, a gene that encodes an extracellular matrix protein (Bisgrove et al., 1991), has strong expression in the rudiment ectoderm (Fig. 2F). We thus annotated the scRNA-seq cluster with strong *Egfi* expression as “Rudiment Ectoderm (Verified)”. To our knowledge, this is the first time that scRNA-seq has been used to capture the transcriptional profile of cells in sea urchin rudiment structures. *Dynlc2*, a gene that encodes a protein component of the dynein light chain (Telmer et al., 2024), is expressed in the tips of the primary podia (Fig. 2G), which may be involved in the intracellular trafficking of vesicles known to be common in this tissue (Burke, 1980). We grouped this with the other rudiment ectoderm clusters. As a final example, *Hox7*, a member of the sea urchin Hox gene complex (Popodi et al., 1996), is expressed in the posterior ectoderm as well as part of the adult rudiment (Fig. 2H) that is transcriptionally distinct from other ectodermal domains (annotated as “Posterior Ectoderm”). Overall, our scRNA-seq atlas of *He* development is a valuable resource for understanding the transcriptional profiles of tissues in late sea urchin larvae.

### Larval cell types in He follow novel developmental trajectories compared to those in species with planktotrophic development

Several *He* larval cell types have different developmental trajectories compared to what was observed in single cell atlases of planktotrophs (Foster et al., 2020; Massri et al., 2021; Paganos et al., 2021). A particularly noticeable difference is with the skeletogenic cell (SKC) cluster (Fig. 3A). In development of planktotrophs, there are two populations of SKCs, one for building the larval skeleton that is derived from the primary mesenchyme cells and a later population for building the adult skeleton that differentiates from the secondary mesenchyme cells (SMCs) (Yajima, 2007). A previous study in *He* found that the SKCs are specified much later in larval development (Davidson et al., 2022). Since larval development in *He* is much shorter than for species with planktotrophic development, we wondered if the late emergence of SKCs may be an indication that the two populations of SKCs are specified differently. To assess this, we first examined the expression of larval skeletogenic genes. Gao and Davidson (2008) established that *Tbr*, *Tel*, *FoxO*, and *FoxB* are expressed in larval, but not adult, SKCs. In the *He* scRNA-seq atlas, few cells in the skeletogenic lineage show co-expression between *Alx1* and *Tbr* (2/106 = 1.8% of *Alx1+* cells in the skeletogenic lineage), *Alx1* and *FoxB* (0/106 = 0% of *Alx1+* cells), and *Alx1* and *FoxO* (7/106 = 6.6% of *Alx1+* cells) and only show moderate co-expression between *Alx1* and *Tel* (24/106 = 22.6% of *Alx1+* cells) (Fig. 3C-F). In contrast, co-expression of these gene pairs is more widespread in our previously published scRNA-seq atlas for *Lytechinus variegatus* (*Lv*), another sea urchin with planktotrophic development (Fig. 3B, C’-F’) (Massri et al., 2021). Nearly all SKCs in *Lv* co-express *Alx1* and *Tbr* (1961/2211 = 88.9% of *Alx1*+ cells), and significant fractions of SKCs co-express *Alx1* and *FoxB* (636/2211 = 28.8% of *Alx1*+ cells), *Alx1* and *FoxO* (517/2211 = 23.4% of *Alx1*+ cells), and *Alx1* and *Tel* (913/2211 = 41.3% of *Alx1*+ cells). These match findings from the analysis of the 6 to 30 hpf scRNA-seq dataset for *He* (Massri et al., 2024) and suggest that a portion of the larval skeletogenic gene regulatory network is no longer functional in *He*.

**Figure 3:**
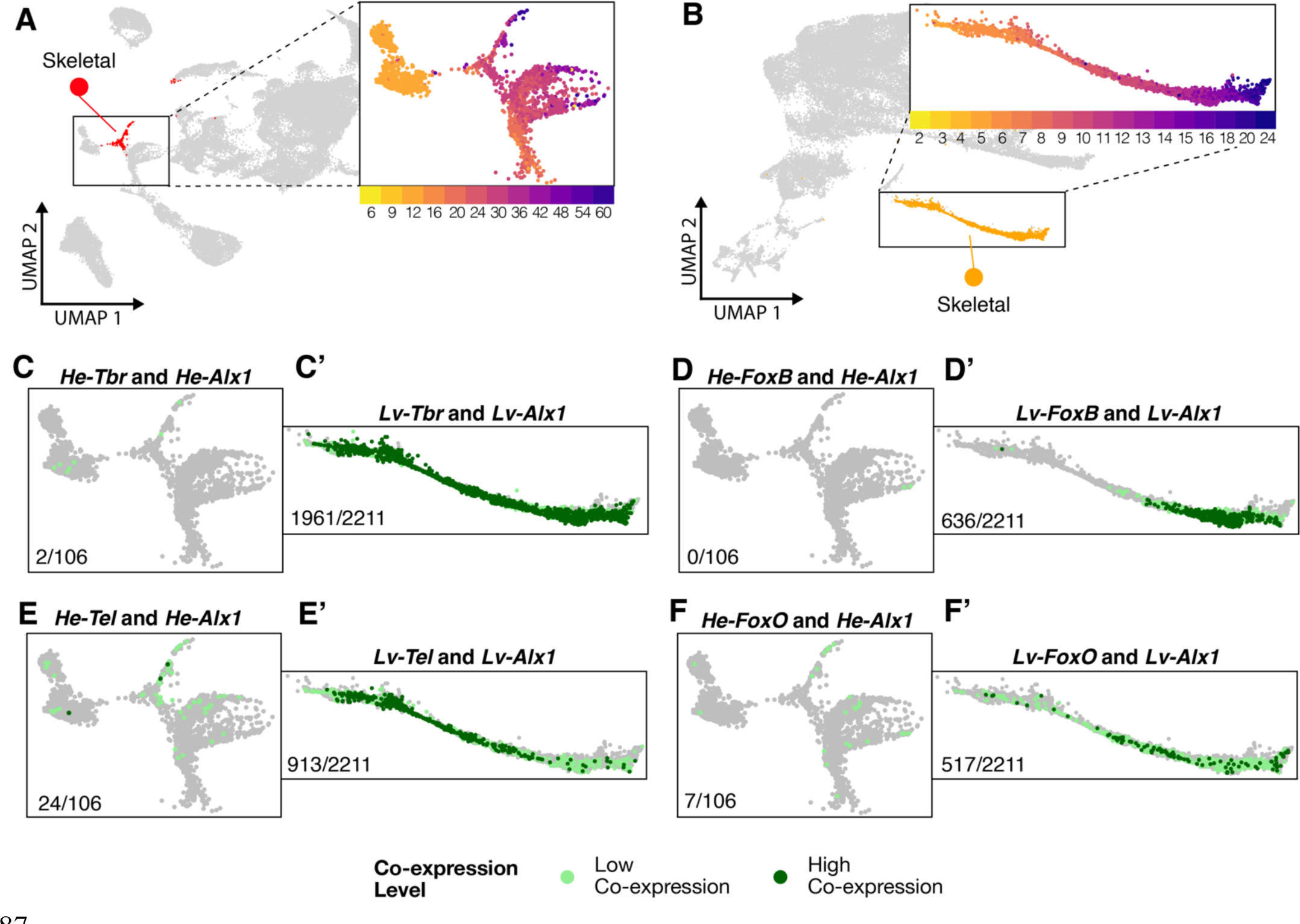
***He* skeletal cells lack expression of genes known to be expressed in the larval SKCs of planktotrophs.** (A) UMAP of *He* development highlighting the SKC cluster (red, background diagram), with an inset of the SKC region of the UMAP colored by time point. (B) UMAP of *Lv* development (retrieved from Massri et al., 2021) highlighting the SKC cluster (orange, background diagram), with an inset of the SKC region of the UMAP colored by time point. (C-F) Cells in the *He* scRNA-seq dataset (zoomed into the SKC region of the UMAP) showing co-expression of *Alx1* and *Tbr* (C), *Alx1* and *FoxB* (D), *Alx1* and *Tel* (E), and *Alx1* and *FoxO* (F). (C’-F’) Cells in the *Lv* scRNA-seq dataset (zoomed into the SKC region of the UMAP) showing co-expression of *Alx1* and *Tbr* (C’), *Alx1* and *FoxB* (D’), *Alx1* and *Tel* (E’), and *Alx1* and *FoxO* (F’). Light green dots represent cells with low co-expression levels, while dark green dots represent cells with high co-expression levels. Numbers at the bottom left of each diagram indicate the number of both the low and high co-expressing cells for each gene pair out of the total number of *Alx1* expressing cells in the SKC lineage for each species.

We then examined whether the larval SKCs in *He* may have acquired characteristics of the adult SKC population in species with planktotrophic development. We focused on the expression of *Scl*, *Ese*, and *GataC*, which are all markers for oral SMCs in planktotrophs (Materna et al., 2013). SKCs in the *He* scRNA-seq dataset show significant co-expression between *Alx1* and *Scl* (93/106 = 87.7% of *Alx1+* cells) and moderate co-expression between *Alx1* and *GataC* (37/106 = 34.9% of *Alx1+* cells) and *Alx1* and *Ese* (19/106 = 17.9% of *Alx1+* cells) (Fig. 4A-C). Double HCRs for *Alx1* and *Scl* in early (36 hpf) *He* larvae provide additional verification that this co-expression event is taking place (Fig. 4D). In *Lv*, co-expression between *Alx1* and oral SMC markers is more limited (Fig. 4A’-C’). The difference is particularly stark for *Alx1* and *Scl*, as only 14.6% (323/2211) of *Alx1+* cells showed co-expression (Fig. 4A’). In addition, while co-expression between *Alx1* and *Scl* begins as soon as *Alx1* starts being expressed in *He* (Fig. 4E), *Scl* expression begins later in *Alx1* expressing cells in *Lv* (Fig. 4F). There were also lower co-expression levels of *Alx1* and *Ese* and *Alx1* and *GataC* in *Lv* (Fig. 4B’-C’). Thus, it appears that SKCs in *He* have gained expression of a subset of genes known to pattern a different mesenchymal population in urchins with planktotrophic development.

**Figure 4:**
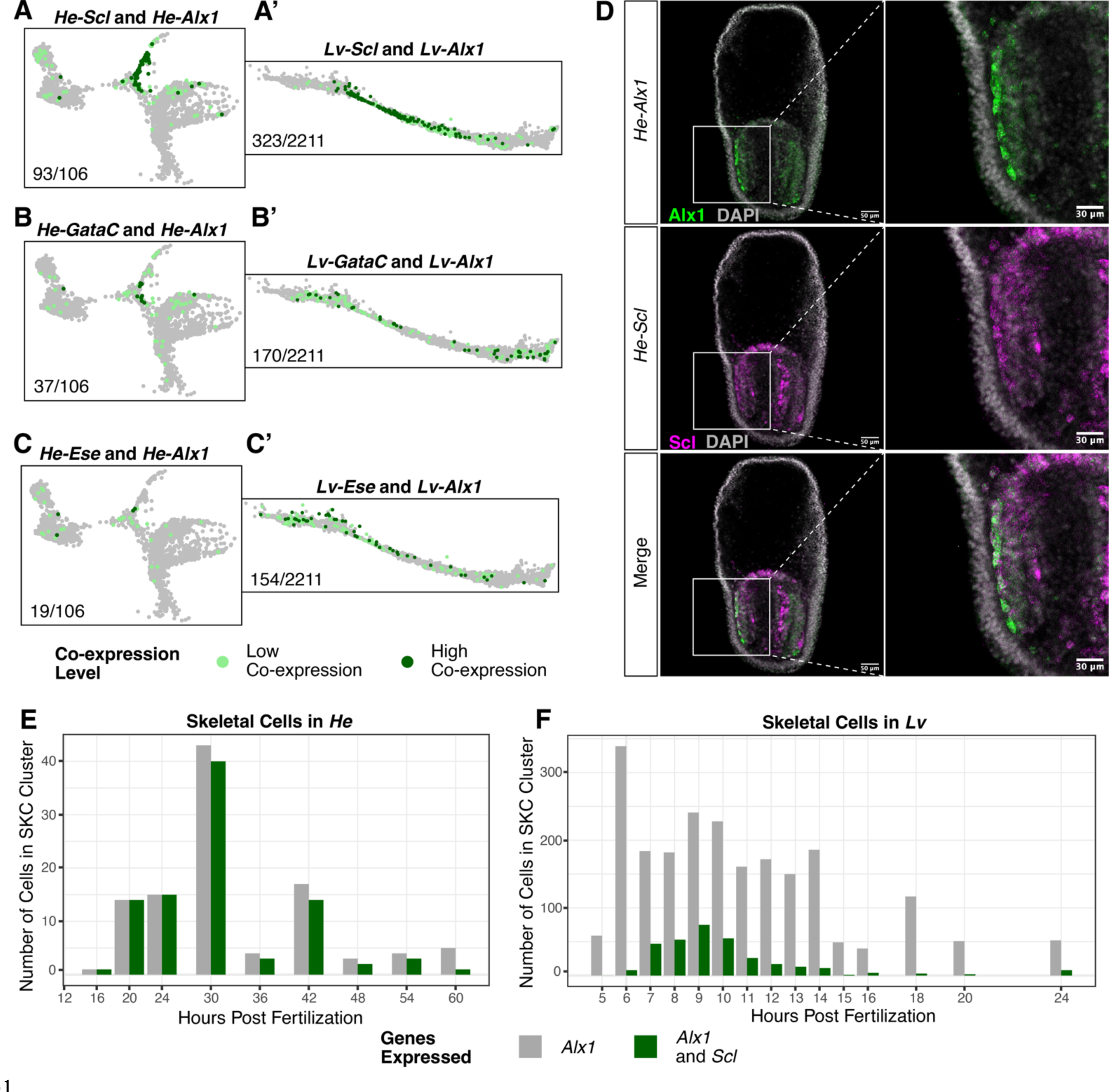
*He* skeletal cells gained co-expression of genes known to be expressed in the secondary mesenchyme cells of planktotrophic sea urchins. (A-C) Cells in the *He* scRNA-seq dataset (zoomed into the SKC region of the UMAP) showing co-expression of *Alx1* and *Scl* (A), *Alx1* and *GataC* (B), and *Alx1* and *Ese* (C). (A’-C’) Cells in the *Lv* scRNA-seq dataset (zoomed into the SKC region of the UMAP) showing co-expression of *Alx1* and *Scl* (A’), *Alx1* and *GataC* (B’), and *Alx1* and *Ese* (C’). Light green dots represent cells with low co-expression levels, while dark green dots represent cells with high co-expression levels. Numbers at the bottom left of each diagram indicate the number of both the low and high co-expressing cells for each gene pair out of the total number of *Alx1* expressing cells in the SKC lineage for each species. (D) HCR micrographs of *Alx1* (green) and *Scl* (magenta) in 36 hpf *He* larvae. Co-expression occurs in a large number of mesenchymal cells surrounding the vestigial gut of early larvae. (E) Chart showing the number of SKCs in the *He* scRNA-seq time course showing expression of *Alx1* (gray) and *Alx1* and *Scl* (green) at each sample time point. (F) A chart showing the same information for the *Lv* scRNA-seq time course.

Another novel larval cell population in the scRNA-seq atlas is a persistent cluster of undifferentiated cells (Fig. 5A). While many cells in this cluster are from the 6 hpf time point, the cluster contains cells from all time points except the last one, 60 hpf (Fig. 5B). This cell population persists throughout larval development, suggesting it is not an artifact of the sample preparation for a single time point. To get a better sense of the gene expression profiles of cells in this cluster, we identified the genes that were significantly enriched in this cluster compared to all others (n = 342) and performed gene ontology (GO) over representation analysis for genes in this list to identify the biological processes that they likely facilitate. None of the enriched genes in this cluster are known marker genes for differentiated cell types in species with planktotrophic development (Table S1). Similarly, most of the enriched GO terms are for categories related to regulation of the cell cycle and the cytoskeleton (Fig. 5D), which are not unique to any differentiated cell type. The undifferentiated nature of cells in this cluster is further demonstrated by how it groups closest to the early ectoderm cluster, which contains multipotent cells from the 6 hpf timepoint, in a phylogenetic tree of cell types (Fig. S3). We then implemented the Waddington-OT trajectory inference algorithm using a previously published pipeline (Massri et al., 2021; Schiebinger et al., 2019) to track cell fate transitions in the dataset. It appears that cells beginning in this cluster transition into several other cell types over the 6 to 60 hpf period covered by the scRNA-seq atlas, especially the ciliary band and larval ectoderm (Fig. 5C). Taken together, these data indicate that the cells in this population start out with undifferentiated transcriptional profiles but eventually differentiate into other cell fates during development. It is important to note that this cluster is unlikely to be the germline due to the absence of co-expression between *Nanos2*, *Vasa*, and *Seawi*, which are known markers of this cell type in planktotrophs (Fig. 5E) (Juliano et al., 2006).

**Figure 5:**
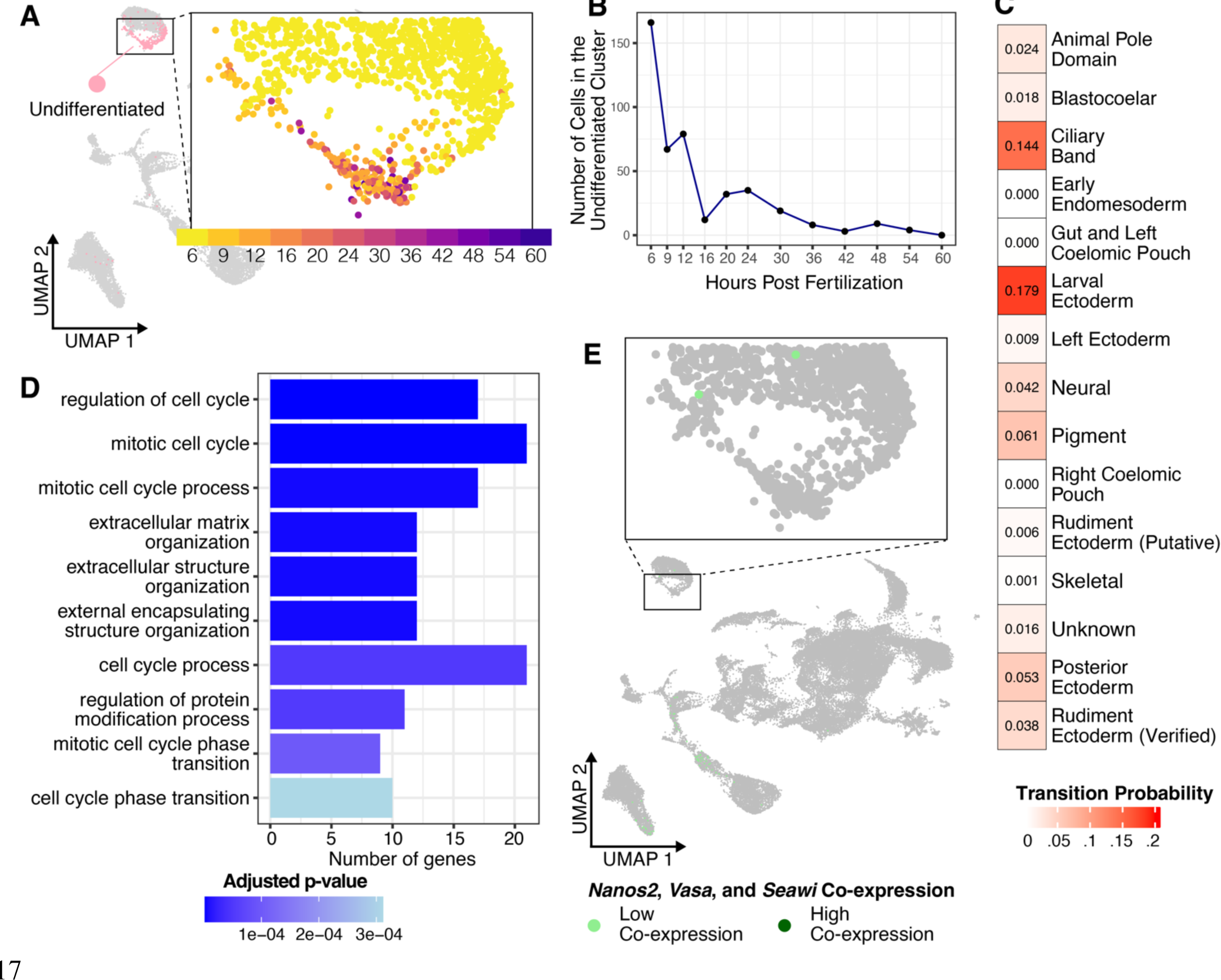
A population of undifferentiated cells persists through the accelerated larval period of *He*. (A) UMAP of *He* development highlighting the undifferentiated cell cluster (pink, background diagram), with an inset of the undifferentiated cell region of the UMAP colored by time point. (B) Plot showing the number of cells belonging to the undifferentiated cell cluster at each time point in the scRNA-seq time course. (C) Transition table showing the Waddington-OT trajectory inference results for the undifferentiated cell cluster. Transition probability values indicate the likelihood that cells beginning in the undifferentiated cell cluster transition into a different cell type identity. (D) Chart showing the top enriched GO terms in the undifferentiated cluster, colored by adjusted p-value. (E) UMAP of *He* development showing co-expression between *Nanos*, *Seawi* and *Vasa*, three of the main markers of the germline in planktotrophic sea urchins.

Finally, the scRNA-seq atlas is a rich resource for understanding the larval nervous system of sea urchins. A previous scRNA-seq study (Paganos et al., 2021) catalogued neural cell types in the early planktotrophic larvae of *Strongylocentrotus purpuratus*. The slower larval developmental process of this species meant that the nervous system was still in the early stages of development at the assayed time point (72 hpf). In contrast, the rapid larval development of *He* meant that the 60 hpf time point of the scRNA-seq atlas was likely to capture a wide array of differentiated neural cell types. To explore this further, we isolated the scRNA-seq clusters expressing pan-neuronal genes (including *Syn4* and *Secrtag*) and performed additional clustering to identify a finer range of neural cell types (Burke, Osborne, et al., 2006). We then annotated each neural cell cluster based on the expression of marker genes identified in previous studies (Fig. S4; Table S2). In total, we identified 15 unique neural cell clusters, including both progenitor and terminally differentiated populations (Fig. 6A-B). Several of these clusters align with previously studied larval neural cell types in sea urchins. Neural progenitor cells from the earliest time points (12 to 20 hpf) show expression of early anterior neuroectoderm (ANE) genes such as *Hbn* and *SoxC*. Differentiated larval neural cell types appear at later stages, including serotonergic neurons expressing *Tph* and postoral and ciliary band neurons expressing *Th*. However, several clusters did not match the expression profiles of known larval neural cell types. The most likely explanation for this was that these were neurons from the developing adult nervous system, given that many adult structures are already present in the rudiments of 60 hpf *He* larvae. We were only able to provide rough annotations for these clusters based on the limited research previously conducted on adult sea urchin nervous systems. For example, the expression of *Isl*, *Pax6*, and *Opsin2* in a late stage cluster matched a previous study that profiled the light receptor neurons found in the podia of adult sea urchins (annotated as “Podia Neurons”) (Ullrich-Luter et al., 2011).

**Figure 6:**
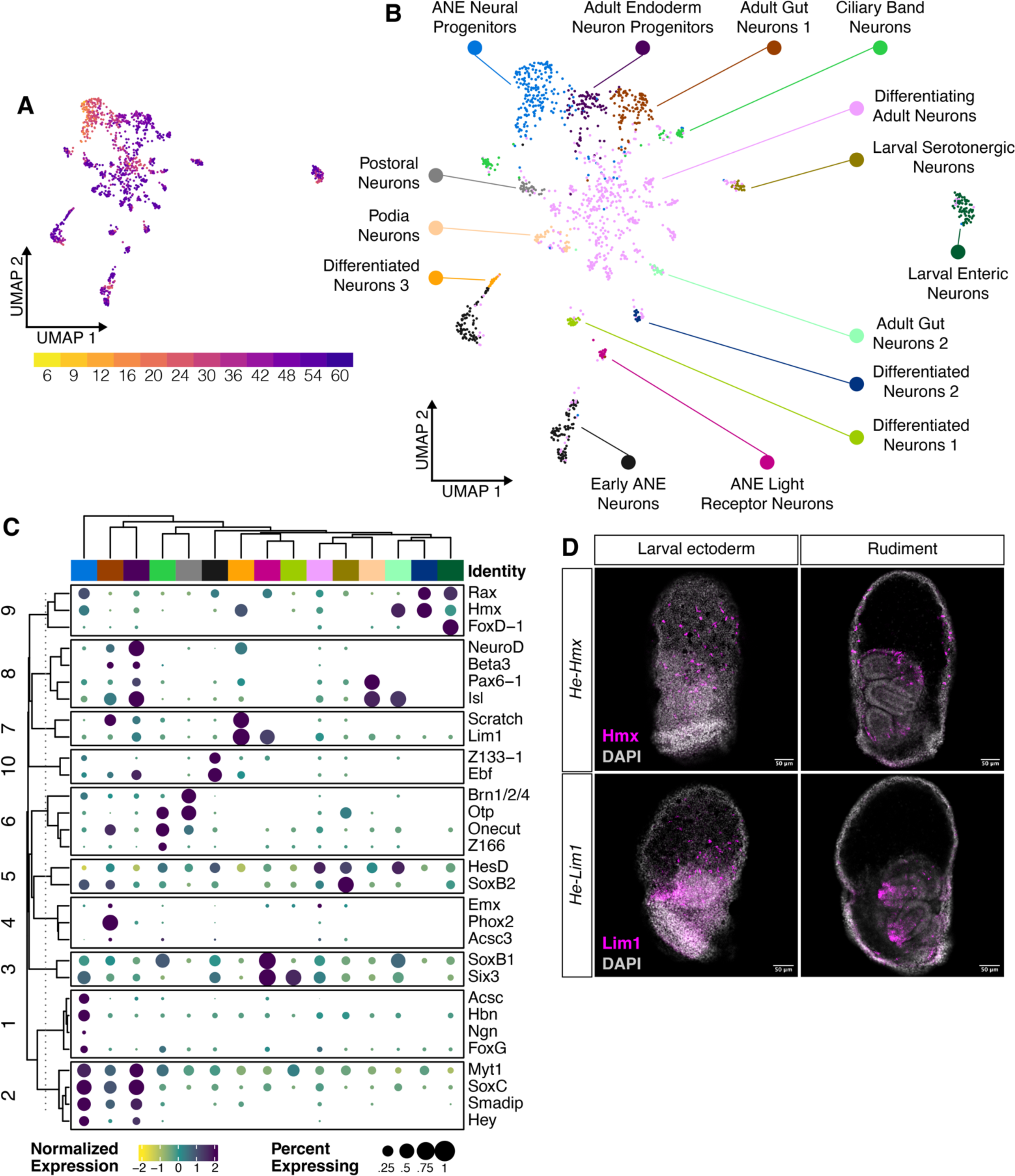
*He* larvae retain ancestral sea urchin larval neural cell types as well as acquire the expression of several putative adult neural cell types. **(A) UMAP of the sub-clustered** neural cells, colored by sample time point. (B) UMAP of the sub-clustered neural cells, colored by neural cell type identities that were assigned based on expression of known larval and adult sea urchin neural patterning genes. Note that several clusters are labeled only as “differentiated neurons”, since they matched no known neural cell type signature. (C) Clustered dot plot of the expression levels of 30 neural TFs in each of the *He* neural cell types. Clustering was performed using k-means clustering with k = 10. (D) HCR micrographs in 56 hpf *He* larvae for *Hmx* and *Lim1*, two transcription factors that are expressed in neurons in *He*. The first column depicts expression in neurons in the larval ectoderm. The second column shows a more interior view of the larva, where these genes have expression in the adult rudiment.

We next turned our attention to the regulation of the neural cell type diversification process in *He*. We compiled a list of 68 transcription factors (TFs) known to be expressed in sea urchin neural cell types (Burke, Angerer, et al., 2006) and selected the 30 TFs with the highest expression levels across all cells in the neural scRNA-seq cluster. We then used k-means clustering to group these genes based on similar expression levels in each of the 15 different neural cell clusters (Fig. 6C). The 10 groups of TFs can be viewed as potential gene regulatory suites controlling the differentiation of neural cell types. Groups 1 and 2 contain TFs with high expression in both ANE and adult neural progenitor cells, such as *Myt1*, *SoxC*, and *Smadip*.

Group 6, with *Brn1/2/4*, *Otp*, *Onecut*, and *Z166*, has high expression in ciliary band and postoral neurons, aligning with studies that found a shared embryonic origin for these neural cell types (reviewed in McClay, 2022). This analysis also identified potential genes that may regulate the differentiation of the unidentified adult neural cell clusters, with the TFs in groups 9 and 10 showing high expression levels in these cells. We noticed that two of these genes, *Hmx* and *Lim1*, also have non-neural expression domains in adult rudiment clusters in the full scRNA-seq atlas. This finding was confirmed using HCR in 56 hpf *He* larvae. Both genes show expression in ectodermal cells with similar morphology to neurons (Fig. 6D, top row) while simultaneously showing expression in rudiment tissues (Fig. 6D, bottom row). It appears that there is overlap between the patterning of the nervous system and the adult rudiment in *He* larvae.

### Different gene regulatory suites appear to control the development of the larval and adult body plans

In organisms with biphasic lifecycles, the switch between the larval and adult body plans does not entirely occur during the brief timeframe of metamorphosis (Formery & Lowe, 2023). Rather, many adult tissues are constructed during larval development, as is the case with the sea urchin rudiment. We thus turned to the *He* scRNA-seq atlas to compare gene expression patterns between early, embryonic stages and later, larval stages when the rudiment is emerging. We began by performing a pseudobulk analysis of the scRNA-seq dataset. We performed fuzzy c-means clustering to group genes with similar temporal expression profiles into 9 clusters (Fig. 7A). Genes that belong to each cluster will have similar shapes to their gene expression profiles over time. Clusters 3, 5, and 8 form a group that shows highest expression during the 6 and 9 hpf time points, which we labeled as “High Early”. Clusters 1, 6, and 7 show highest expression in the middle of the time course from 12 to 30 hpf, which we labeled as “High Middle”. Finally, clusters 2, 4, and 9 peak in expression at 36 hpf or later, which we labeled as “High Late”. We then performed GO over representation analysis to see whether certain biological processes were enriched for genes in some clusters compared to others (Fig. 7B). “High Early” clusters are enriched for functions related to metabolism, cell cycle regulation, and organelle remodeling, which may be due to the rapid cell divisions that occur early in embryogenesis. Many of the enriched terms in the “High Middle” clusters are for cell division related processes, as well as transcription and translation. This may reflect widespread larval cell differentiation during late embryogenesis. Clusters in the “High Late” category show enrichment for nervous system functions, aligning with the rapid diversification of neural cells during later larval stages (Fig. 6). There is also enrichment for terms related to cell migration and movement in the “High Late” category. Cell movement may be common in later developmental stages as larval cells are rearranged during the morphogenesis of adult tissues.

**Figure 7:**
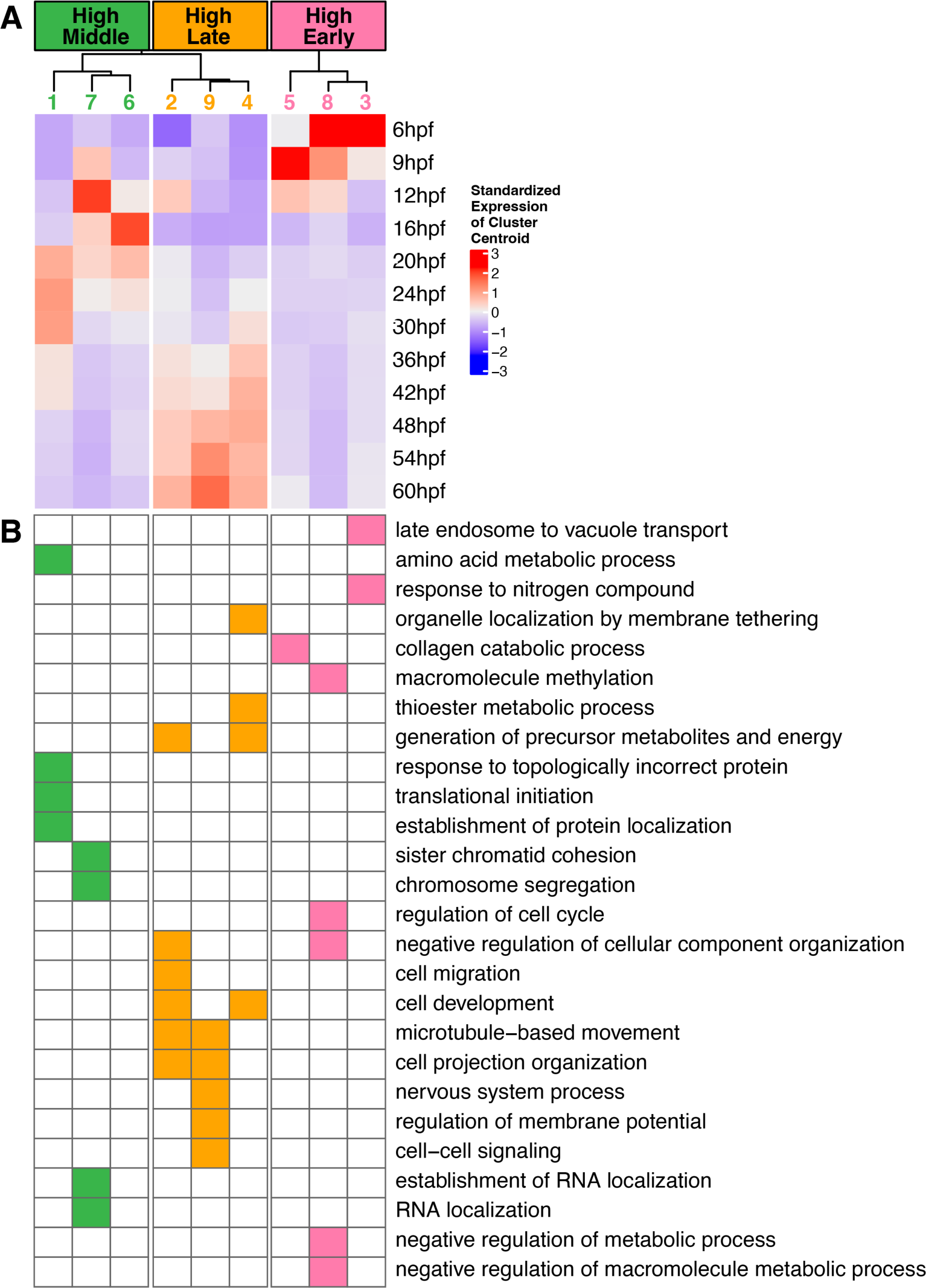
Genes with late expression peaks in the scRNA-seq time course appear to participate in a unique set of biological processes. (A) Heatmap showing the centroids of expression profiles from the Mfuzz analysis of pseudobulked scRNA-seq data at each time point. (B) Heatmap showing gene ontology term enrichments for each of the Mfuzz cluster profiles. A filled square indicates that there was a statistically significant enrichment of genes that perform the indicated biological process in the Mfuzz cluster.

The gene function analysis suggested that a unique set of processes may be taking place during later stages of larval development in *He*. This prompted us to ask whether the well-defined GRN for sea urchin embryonic development (Davidson et al., 2002; Peter & Davidson, 2011; Rafiq et al., 2012; Sethi et al., 2009; Su et al., 2009) plays a similarly important role in late larval and adult development. Many of the genes in the embryonic GRN are transcription factors and signaling molecules that are commonly used in many developmental processes, so it would not be unexpected to see similar genes being used in both time frames. To assess this question, we retrieved a curated list of embryonic GRN genes used in previous analyses (Davidson et al., 2022) and classified whether the temporal expression profile of each gene fell into the “High Early”, “High Middle”, and “High Late” categories obtained from the fuzzy c-means analysis. As a comparison, we also generated a list of all the potential DNA-binding transcription factors (TFs) that are found in the *He* genome based on GO terms and a previous curated list (see Methods section). The number of embryonic GRN genes and TF genes that fell into each temporal category is shown in Fig. 8A. Note that many, but not all embryonic GRN genes are TFs, so the numbers of GRN genes and general TFs in each category are not directly comparable. Many of the GRN genes were also included in the broader TF category. However, it was striking to see that the vast majority of GRN genes have expression profiles that peak early on or in the middle of *He* development (100 with early or middle peaks vs. 38 with late peaks). In comparison, there are roughly equal numbers of TFs that have expression peaks in each temporal category. It thus appears that genes in the embryonic GRN may play a reduced role in patterning the late larvae and adult rudiment of *He*. Of the few embryonic GRN genes that have late expression peaks, several have roles in patterning the larval nervous system in planktotrophs, such as *Scratch* and *Isl* (Fig. 8B) (Slota et al., 2019).

**Figure 8:**
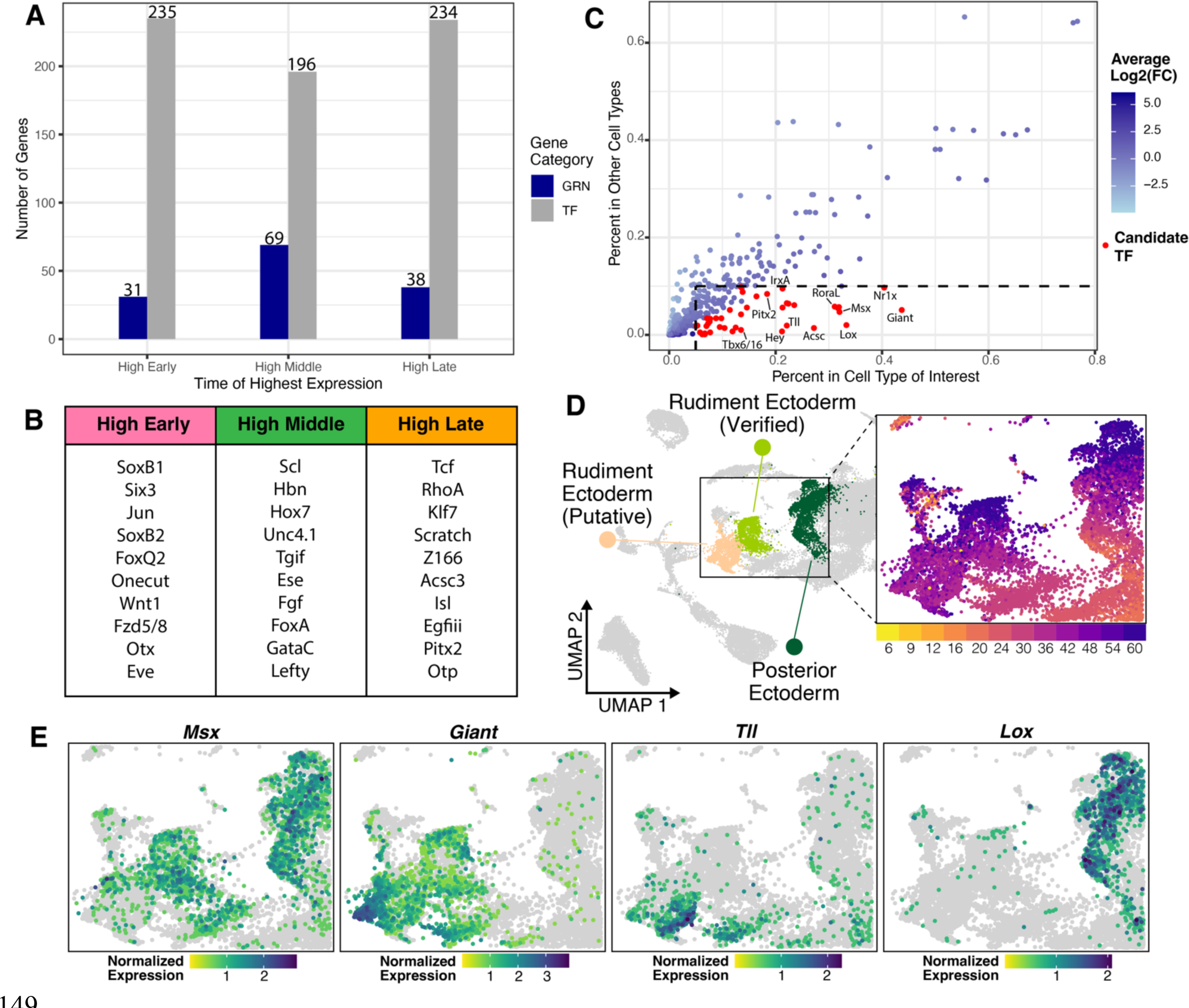
Different gene regulatory suites appear to control the development of the larval and adult body plans. (A) Chart showing the number of TFs (gray) and embryonic GRN genes (blue) with expression peaks in each temporal category. (B) The top 10 expressed embryonic GRN genes with expression peaks in each temporal category. (C) Plot showing the percent of cells showing expression of TFs with late peaks in cell types of interest (rudiment ectoderm and posterior ectoderm) versus all other cell types, colored by their average log2-fold change in the cell types of interest (as a measure of enrichment). Points colored red indicate genes with average log2-fold changes > 0.9 and that were expressed in more than 5% of cells in the cell type of interest and less than 10% of cells in the other cell types. (D) UMAP of *He* development highlighting the adult rudiment clusters. (E) UMAPs of *He* development zoomed in on the adult rudiment clusters, colored by expression levels for TFs potentially controlling adult rudiment morphogenesis.

Since embryonic GRN genes appear to have lower expression levels at later stages, we hypothesized that adult rudiment development in *He* may be controlled by a new set of transcriptional regulators. We moved to generate a list of candidate TFs that could play a regulatory role in this process. We returned to the scRNA-seq dataset and focused on the rudiment ectoderm and posterior ectoderm cell clusters, which are the primary cell types in the scRNA-seq atlas with verified expression patterns in the rudiment (Fig. 8D). Of the 234 TFs that have late expression peaks, we filtered for genes that had high expression in the aforementioned cell types and low expression elsewhere. This resulted in a set of 33 unique TFs (Fig. 8C, red points; Fig. S5; Table S3) that are potentially important regulators of adult rudiment development. Some of these genes are already known to pattern the rudiment in *He* (*Msx*, Fig. 8E) (Koop et al., 2017; Wilson et al., 2005), but others appear to be novel transcriptional regulators of sea urchin development. For example, *Giant* and *Tll* show strong localized expression in the rudiment ectoderm cell type clusters (Fig. 8E). *Lox*, which is known to pattern the hindgut in planktotrophic larvae (Cole et al., 2009), is expressed in the posterior ectoderm, suggesting it may have an additional role in rudiment patterning (Fig. 8E). These 33 genes represent a list of candidate TFs to experimentally manipulate in future studies to understand the regulation of adult rudiment development.

## Discussion

In this paper, we presented a comprehensive scRNA-seq atlas of embryonic and larval development in *He*. Building off the insights gained from previous datasets in sea urchins with planktotrophic development (Foster et al., 2020; Massri et al., 2021; Paganos et al., 2021) and *He* (Massri et al., 2024), the present analysis provides single cell transcriptomic data for late stages of urchin larval development that lacked coverage in prior studies. Research on cell type trajectories in later larval stages has historically been difficult due to the long periods required to raise planktotrophic larvae through metamorphosis. In contrast, the accelerated developmental process of *He* makes it a useful model for studying the full time frame of larval developmental events, as well as the initial stages of adult morphogenesis (Wray, 2022). By using HCRs to map marker genes from the scRNA-seq dataset onto discrete tissues, we present some of the first transcriptomic data that is specific to the adult rudiment. These data are an important resource for studying the fates of larval cell types during adult development and allow us to start uncovering the GRN that controls this process. We also observed two important patterns regarding the biphasic lifecycle of *He*. On one hand, several larval cell fate trajectories have been modified compared to those in planktotrophs. At the same time, the larval developmental program appears distinct from the adult program, potentially allowing for each stage to evolve in a modular fashion.

### Modifications to larval cell type trajectories during the evolution of lecithotrophy

Prior research in *He* embryos revealed that several long-conserved cell fate trajectories have been reprogrammed during the evolution of lecithotrophy (Davidson et al., 2022; Parks et al., 1988; Wray & Raff, 1990). We observed similar trends with larval cell types in our scRNA-seq atlas. First, there was a notable change in the expression profiles of SKCs. In *He*, our co-expression analyses indicate that the SKCs lack expression of larva-specific skeletal genes but have gained expression of genes known to mark SMCs, which are likely the progenitors of the adult SKC population in planktotrophs (Figs. 3 and 4) (Gao & Davidson, 2008; Yajima, 2007). This aligns with research in *He* showing that SKCs are no longer specified early in embryonic development and have lost expression of key components of the ancestral GRN (Davidson et al., 2022; Massri et al., 2024). The most likely explanation is that the ancestral larva-specific SKC population has been lost in *He*. This raises the question of what is the origin of the cell population that patterns *He*’s vestigial larval skeleton. One possibility is that early-arising adult SKCs fulfill this role. A few of the cells that are intended to produce skeletal structures in the adult rudiment may instead respond to leftover patterning cues in the ectoderm that evolved to guide larval SKCs. Another possibility is that the larval skeleton may be produced by SMC-like cells that undergo transfating toward skeletogenic fates. Removal of the SKC progenitors in the embryos of species with planktotrophic larvae is known to trigger a subset of SMCs to activate the downstream SKC GRN (Ettensohn et al., 2007; Ettensohn & McClay, 1988; Sharma & Ettensohn, 2011). *He* embryos may have retained this transfating capability for the purposes of normal development. In either case, the elimination of the need to specify larval SKCs in the early embryo may have facilitated the acceleration of larval development during the evolution of lecithotrophy. Alternatively, the loss of most larval SKCs may be a consequence of the acceleration.

We also reported the emergence of an undifferentiated cell cluster present in the *He* embryo and early larva. It does not appear that these cells are a classical stem cell population, given their failure to express known pluripotency or germline related genes (Fig. 5D; Table S1). We can also rule out that the cluster is the hypothesized “set aside” cell population that is thought to supply most of the cells needed during metamorphosis (Davidson et al., 1995; Peterson et al., 1997), since few cells are present in the cluster at later timepoints (Fig. 5B). A more likely possibility is that these cells represent a novel cell state that is held back from differentiating into terminal cell fates early in development. It is already known that cell fate specification is delayed in *He* relative to species with planktotrophic development (Davidson et al., 2022) even though the overall larval developmental period is accelerated. These cells may be a dedicated population that is reserved for rapid differentiation once initial patterning events take place in the embryo.

This is supported by the fact that the undifferentiated cluster groups with the early ectoderm cluster (which mostly consists of undifferentiated cells from the 6 hpf time point) in the cell type dendrogram (Fig. S3), despite containing cells from several later time points. While the exact fates of cells in this cluster are unknown, there is no analogous cell population that appears in scRNA-seq atlases of planktotrophs (Foster et al., 2020; Massri et al., 2021; Paganos et al., 2021) that arises early in development and persists over multiple time points. This cell state could therefore be a feature unique to lecithotrophs and may be a consequence of altered cell specification patterns in the *He* embryo.

A third larval cell type that showed a notable shift was the neural cells. Work from the past couple of decades has led to a thorough understanding of the specification and functions of the limited set of neurons found in early sea urchin larvae (reviewed in McClay, 2022). Many of these neural cell types are captured by our scRNA-seq atlas, including the larval serotonergic neurons and postoral neurons (Fig. 6). At the same time, we observed the emergence of several putative adult neural cell types at the later developmental time points in the atlas. Early appearance of adult neural cell types is likely associated with the acceleration of larval development in *He*. We also noticed that several transcription factors expressed in the putative adult neurons are expressed in clusters corresponding to adult rudiment tissues, suggesting that the processes of patterning the adult nervous system and broader pentaradial adult body plan are tightly connected (Ferkowicz & Raff, 2001; Sly et al., 2002). This aligns with recent work showing how anterior-posterior patterning genes––which primarily pattern the neuroectoderm in hemichordates and chordates––have been redeployed to pattern the pentaradial adult body plan of sea stars (Formery et al., 2023). Nervous systems and body plans have frequently been observed to evolve hand-in-hand (Holland, 2016; Holland et al., 2015), so further research on the development of the sea urchin adult nervous system could help explain the transition from bilateral to pentaradial symmetry in echinoderm evolution.

### The emergence of a new regulatory network governing adult body plan development in sea urchins

Our data also allow us to address questions about the regulation of adult morphogenesis in sea urchins. It has previously been unclear how organisms with biphasic lifecycles code for two different body plans in the same genome (Formery & Lowe, 2023). Our data suggest that a largely new set of regulatory genes controls the adult developmental process in sea urchins. A key finding is that most genes from the sea urchin embryonic GRN show the highest expression in the embryo and early larvae of *He* and have lower expression during later larval time points (Fig. 8A). To a certain extent, this is not entirely surprising because the GRN was constructed based on early developmental time points (Davidson et al., 2002). However, many of the genes in the embryonic GRN are part of signaling systems used in a diverse array of developmental patterning events in metazoans, such as Wnt/Fzd and TGFß factors (Clevers, 2006; Wu & Hill, 2009), so the reduced role of some of these genes in adult patterning events is notable. This lack of overlap may partially be due to the drastic switch from bilateral to pentaradial symmetry during metamorphosis in echinoderms (Peterson et al., 2000; Smith, 2008). Not all of the genes that pattern the bilateral larva may be suited for patterning the pentaradial adult, requiring the introduction of a new suite of developmental regulators. Thus, the embryonic GRN appears to play a reduced role in regulating adult morphogenesis.

In place of the embryonic GRN, we generated a candidate list of 33 TFs that may fulfill the regulatory requirements of patterning the pentaradial adult rudiment (Fig 8E; Fig. S5; Table S3). Unfortunately, the scRNA-seq expression patterns of these genes alone do not reveal the upstream mechanism responsible for specifying the initial pentaradial pattern. To address this question, we can turn to previous embryological and gene expression studies in *He*. It appears that signaling between endomesodermal tissues and the overlying larval ectoderm is necessary for the invagination of the vestibule (Minsuk et al., 2005; Minsuk & Raff, 2002), and these interactions may also establish pentaradial symmetry in both tissues (Minsuk et al., 2009). Koop et al. (2017) followed up on this work by examining the expression of genes in the Nodal-BMP signaling network in early *He* larvae, which is known to pattern the dorsal-ventral and left-right axes of sea urchin embryos (reviewed in Molina et al., 2013). Of the genes they examined, *Bmp2/4* had the earliest pentaradial expression pattern in the rudiment, suggesting that the Nodal-BMP network could be responsible for the initial establishment of the adult body plan.

This led them to claim that the embryonic GRN may play a major role in patterning the rudiment, contradicting our data. One potential explanation for this discrepancy is that key signaling systems like the Nodal-BMP network play a role in the upstream establishment of pentaradial symmetry, but the downstream transcription factors and effector genes that pattern specific adult tissues may lie outside the scope of the embryonic GRN. Extending scRNA-seq atlases to later larval stages in other sea urchin species, along with targeted perturbation studies, will allow us to construct a GRN specific to sea urchin adult morphogenesis.

### The evolution of biphasic lifecycles

In summary, our analysis is valuable for understanding the switch in developmental strategy that occurred during the evolution of *He*, as well as the process of adult body plan construction in general across sea urchins. By examining a scRNA-seq time course that covers most of larval development in *He*, we were able to assess how larval cell types are established in sea urchins and how these may be involved in the transition to the adult body plan. While our data do not capture gene expression events that occur immediately before and during metamorphosis, this work does provide a major advance to our knowledge of how late larval development is regulated in sea urchins. Furthermore, the fact that sea urchins may use two different GRNs to pattern the larval and adult body plans could address uncertainties about the origins of biphasic lifecycles in bilaterians (Davidson et al., 1995; Nielsen, 2013; Raff, 2008; Valentine & Collins, 2000). Instead of larval patterning mechanisms being extensively co-opted for patterning the adult, or vice versa, the two processes appear distinct. A scenario where the larval or adult phase was secondarily added to the lifecycle by simply co-opting pre-existing regulatory networks (Sly et al., 2003) appears unlikely. Answering this question with more confidence will require comparative studies with bilaterian species in other phyla.

Our results also allow us to start to develop a model for how biphasic lifecycles can evolve. The acceleration of development in sea urchins that have lecithotrophic larvae could mean that adult development in *He* follows different patterns compared to species that have planktotrophic larvae. Even so, the adults of *He* and its sister species *Ht*, which has planktotropic larvae, have remarkably similar morphologies (Byrne & Ohara, 2017). Although it is possible that each species relies on a different developmental program to arrive at its adult form, a more parsimonious explanation is that *He* has retained many of the mechanisms found in species with planktotrophic development. At the same time, the divergence of larval patterning strategies in *He* compared to planktotrophs indicates that under the appropriate selective conditions, developmental processes can be highly labile. We hypothesize that the larval and adult portions of biphasic lifecycles can be viewed as semi-independent modules, given that they may be controlled by largely distinct sets of regulatory factors. Taken together, these patterns suggest that the simultaneous conservation and modification of different stages of biphasic lifecycles may have contributed to the diversity of metazoan developmental strategies seen today.

## Materials and Methods

### Single cell RNA-sequencing analysis

#### Data retrieval

*He* embryos and larvae raised at 23°C were sampled at 12 time points from 6 to 60 hpf (6, 9, 12, 16, 20, 24, 30, 36, 42, 48, 54, and 60 hpf) to yield scRNA-seq libraries. All libraries were prepared and sequenced during data collection for the work described in Massri et al. (2024).

However, only samples from the 6 to 30 hpf time points were used in the previous analysis. For the present analysis, data for the same 6 to 30 hpf time points, as well as five additional time points up to 60 hpf, were analyzed with a focus on larval and adult rudiment development. Gene expression counts tables that result from processing the sequencing data using Cell Ranger (Zheng et al., 2017) were retrieved for all 12 time points and were used as input into downstream analysis steps.

#### Data filtering and normalization

After obtaining the counts tables, the R package Seurat v4.3.0 was used for downstream filtering and normalization steps (Hao et al., 2021). The counts tables for each time point were merged into the same Seurat object, which was filtered for high-quality cells with nFeature_RNA > 200, nCount_RNA < 10000, and nFeature_RNA < 4000. In addition, a new column “percent.Rb” was added to the metadata to identify the proportion of transcripts in each cell that have names matching the regular expression pattern “\\b\\w*Rp[sl]\\w*\\b”, which identifies genes that encode ribosomal proteins. The function SCTransform was used with the“glmGamPoi” method and 6000 variable features to normalize, rescale, and identify variable features in the dataset (Choudhary & Satija, 2022). The parameter vars.to.regress = “percent.Rb” was also passed to the SCTransform function to regress out expression signals from ribosomal proteins.

#### Dimensionality reduction, clustering, and visualization

The RunPCA function was then used to perform principal component analysis and dimensional reduction on the Seurat object, using 200 principal components total. Clustering of different cell types was then performed using the FindNeighbors function with the first 195 principal component dimensions followed by running the FindClusters function with resolution = 3. This detected 57 distinct cell clusters across all 12 sample time points, which was a number that remained relatively robust even at higher resolution values. Each of the 57 cell clusters was assigned a cell type identity based on expression of known sea urchin developmental GRN genes and *in situ* hybridization patterns (Fig. 2C-H; Fig. S2; Table S1). To visualize the cell clusters in two-dimensional space, the uniform manifold approximation and projection (UMAP) algorithm was applied using the RunUMAP function. Unless otherwise noted, all gene expression UMAPs were generated using the FeaturePlot function from Seurat or the FeaturePlot_scCustom wrapper function from the R package scCustomize v2.0.1 (Marsh et al., 2023).

#### Neural cell analysis

The clusters that were annotated as “Neural” in the full Seurat object were isolated into a separate Seurat object using the subset function. The neural-specific object was re-clustered using the FindNeighbors function with the first 195 principal component dimensions and the FindClusters function with resolution = 0.8. This resulted in the identification of 17 neural cell types, which was a number that remained relatively robust to increases in the resolution value. Published data on neural marker genes was used to assign each of these clusters a neural cell type identity (Fig. S4; Table S2). Finally, the UMAP algorithm was applied to the neural-specific object using RunUMAP to re-project the neural cells into UMAP space.

To evaluate the transcription factors used to pattern neural cell types, the list of neural transcription factors identified in the *Strongylocentrotus purpuratus* genome was retrieved from Burke, Angerer, et al. (2006). The orthologues of these genes in the *He* genome were identified using the annotations published in Davidson et al. (2022). The top 30 expressed neural transcription factors were selected based on highest average expression levels across all the neural cell types. The expression levels of these genes in each neural cell type were plotted in Fig. 5C using the Clustered_DotPlot function from scCustomize. K-means clustering was used to group the expression patterns of these genes in neural cells into 10 clusters.

#### Co-expression analyses for skeletal and set aside genes

The Seurat function WhichCells was used to highlight cells in atlas that co-expressed two different genes of interest. Cells with expression levels greater than 0.5 for both genes were designated as “low co-expression”, while cells with expression levels greater than 1 for both genes were designated as “high co-expression”. Co-expressing cells were plotted in two-dimensional UMAP space.

scRNA-seq data for *Lytechinus variegatus* (*Lv*) embryos and larvae, a species with planktotrophic development, was retrieved from the scRNA-seq atlas presented in Massri et al. (2021). This atlas contains hourly or bi-hourly scRNA-seq timepoints of development in *Lv* from 2 to 24 hpf. The same steps used for analyzing SKC co-expression patterns in the *He* scRNA-seq atlas were applied to the *Lv* scRNA-seq atlas.

#### Waddington-OT trajectory inference

Waddington-OT trajectory inference (v1.0.8) was performed as previously described (Massri et al., 2024; Massri et al., 2021; Schiebinger et al., 2019), with only a few minor modifications for the full 6-60 hpf time course. Briefly, the full Seurat object was used to generate a SCTransform normalized expression matrix showing gene expression for each cell. Separate files with cell type annotation and UMAP embedding assignments for each cell were also generated. Cell division rates for each cell type were estimated based on the expected number of cells that are supposed to be present at each developmental time point. These rates along with the expression matrix were used to compute transport maps between each of the time points, with parameters set at epsilon = 0.05, lambda1 = 1, lambda2 = 50, and growth_iterations = 20. The transition_table function was used to generate the plot seen in Fig. 5C.

#### Pseudobulk analysis and gene expression profile clustering

To perform pseudobulking of the scRNA-seq data, raw gene expression count data was extracted from the Seurat object. Gene expression counts were aggregated across the cells in each time point, resulting in a matrix with the 12 time points as columns and genes as rows. The counts table was input into the DESeq function from the R package DESeq2 v1.38.3 (Love et al., 2014) to normalize the gene expression data using the median of ratios method. The normalized counts data were then exported from the DESeq object for input into the Mfuzz pipeline.

The R package Mfuzz v2.58.0 (Futschik & Carlisle, 2005; Kumar & Futschik, 2007) was used to perform fuzzy c-means clustering of temporal gene expression profiles. First, genes with normalized expression counts less than 5 at any time point were filtered out of the dataset. The data was filtered to exclude genes with NA expression values and then standardized (for a total of 19585 genes). The fuzzifier parameter m was estimated using the mestimate function. Finally, the normalized and standardized counts data, along with the estimate for m, were provided to the mfuzz function to cluster the gene expression profiles. The optimal number of clusters (value of c) was determined as described in Israel et al. (2016). Briefly, the numb er of clusters was chosen to be as high as possible without having the correlation between any pair of cluster centroids exceed 0.85. This was to ensure that the data was not over clustered and that each cluster showed a relatively distinct pattern from the others. Nine clusters (c = 9) was selected as the optimal value. The centroids of each cluster were plotted (Fig. 6A) using the R package ComplexHeatmap v2.18.0 (Gu et al., 2016). This package was also used to perform k-means clustering on the Mfuzz cluster profiles based on Euclidean distance with k = 3. The first group contained Mfuzz clusters 3, 5, and 8, which were designated as “High Early” because their member genes peaked in expression before 12 hpf. The second group contained Mfuzz clusters 1, 6, and 7, which were designated as “High Middle” because their member genes peaked in expression between 12 and 36 hpf. The third group contained Mfuzz clusters 2, 4, and 9, which were designated as “High Late” because their member genes peaked in expression after 36 hpf.

#### Protein functional annotation and gene ontology analysis

The Linux package InterProScan v5.64-96.0 (Jones et al., 2014) was used to assign functional annotations to the protein coding sequences in the *He* genome. Gene models from *He* were retrieved from Davidson et al. (2022) and translated into peptide sequences. The main InterProScan analysis pipeline was run on these peptide sequences to assign GO terms using all the functional annotation databases contained in the standard InterProScan distribution.

Gene ontology over representation analysis was performed using the R package clusterProfiler v4.10.0 (Wu et al., 2021). The compareCluster function was used to compare overrepresented GO terms between genes assigned to each of the 9 Mfuzz clusters. “Biological Process” GO terms were specifically analyzed, with pvalueCutoff = 0.05, pAdjustMethod = “BH”, and qvalueCutoff = 0.2. The pairwise_termsim and treeplot functions from the R package enrichplot v1.22.0 (Yu, 2023) were used to plot the heatmap in Fig. 6D.

Gene ontology over representation analysis was also used to identify the top “Biological Process” GO terms for genes enriched in the undifferentiated cell cluster. The Seurat FindMarkers function was used to identify identify the genes significantly enriched in this cluster (n = 342), with m.pct = 0.25, logfc.threshold = 0.25, and p_value_adj < 0.05. The enrichGO function from clusterProfiler was run to compare these genes against all other genes in the genome as a background set, with pvalueCutoff = 0.05, pAdjustMethod = “BH”, and qvalueCutoff = 0.2.

#### Embryonic gene regulatory network analysis

A list of genes in the ancestral sea urchin embryonic GRN (192 total) was retrieved from Davidson et al. (2022). In order to access gene expression information across all analyses, only embryonic GRN genes that were retained in the Mfuzz and Seurat analyses were analyzed (138 total). Based on the Mfuzz cluster to which each gene was assigned, embryonic GRN genes were classified as belonging to the “High Early”, “High Middle”, or “High Late” expression profile groups, as described above.

#### Transcription factor analysis

When assembling a list of all the predicted TFs in the *He* genome, the goal was to evaluate as many genes as possible, rather than relying on a restricted set of verified genes. Thus, lists of predicted TFs from two sources were combined along with a list of zinc finger proteins, which are frequently transcription regulators (Klug, 2010). First, the list of TFs and zinc finger proteins used in a previous genomic analysis of *He* (Devens et al., 2023) was retrieved. This was combined with a larger list of TFs predicted from their associated gene ontology terms. Genes with the following terms were included: GO:0003700 (“DNA-binding transcription factor activity”), GO:0001216 (“DNA-binding transcription activator activity”), GO:0001227 (“DNA-binding transcription repressor activity, RNA polymerase II-specific”), GO:0001228 (“DNA-binding transcription activator activity, RNA polymerase II-specific”), GO:0000981 (“DNA-binding transcription factor activity, RNA polymerase II-specific”), and GO:0001217 (“DNA-binding transcription repressor activity”) (Gene Ontology et al., 2023). In total, the combination of these two lists resulted in 1400 transcripts in the *He* genome that are predicted to be TFs, some of which may be duplicated copies of the same gene. Only TFs that were retained in the Mfuzz analysis (n = 665) were used for downstream steps.

To identify the key TFs responsible for patterning adult rudiment structures in *He*, the TF list was filtered for genes that had temporal expression profiles belonging to the “High Late” category (n = 234). The Seurat function FoldChange was used to identify genes from this list that were enriched in the “Rudiment Ectoderm (Verified)”, “Rudiment Ectoderm (Putative)”, and “Posterior Ectoderm” clusters in the scRNA-seq atlas. TF genes were retained that had pct.2 < 0.1, pct.1 > 0.05, and avg_log2FC > 0.9 in any one of these cell type categories. This resulted in a list of 33 unique genes that showed relatively high, enriched expression in adult tissues in *He* larvae (Fig 8E; Fig. S5; Table S3).

All computational analyses were conducted using R v4.2.0 or v4.3.0 (depending on package compatibility) and Python v3.8.

### *In situ* gene expression analysis

#### Embryo fixation

*He* embryos and larvae from desired developmental stages were fixed overnight in 4% paraformaldehyde in artificial sea water (ASW) at 4°C. Embryos were washed once with ASW and once with ice-cold methanol. These were then moved to a fresh aliquot of methanol and stored at -20°C until used for later steps.

#### Hybridization chain reaction probe design

DNA probes for hybridization chain reaction *in situ*s were designed by Molecular Instruments or using the insitu_probe_generator python script from Kuehn et al. (2022). For the non-Molecular Instruments probes, the script selected the maximum number of probes to hybridize with the exon sequence of the gene of interest without overlapping, and the probes were designed with attachments for either B1, B2, or B3 hairpins for the HCR amplification step. Probes were synthesized using the oPools Oligo Pools service from Integrated DNA Technologies. Upon receipt, the probes were diluted to 1 µM in RNAse-free water and stored at -20 °C. See Table S4 for the probe sequences.

#### Hybridization chain reaction in situs

Whole mount fluorescent *in situ* hybridization was performed on fixed *He* embryos and larvae using the v3.0 hybridization chain reaction method, following the sea urchin-specific protocol published in Choi et al. (2016) with some modifications. Unless otherwise noted, all washes were performed at room temperature. Fixed samples were rehydrated stepwise from methanol into PTw (1x PBS and 0.1% Tween 20). To permeabilize the embryo membranes, the samples were incubated for 30 minutes in a detergent solution containing 1% SDS, 0.5% Tween 20, 50 mM Tris-HCl, 1 mM EDTA, and 150 mM NaCl. Samples were washed in PTw, and a post-fixation step was performed by incubating the embryos in 4% paraformaldehyde in PTw for 25 minutes. The fixative was removed with five washes of PTw.

The samples were then pre-incubated in Probe Hybridization Buffer (Molecular Instruments) for 30 minutes at 37°C. Probe solutions were prepared by adding 2 µL each of DNA probes for one or two genes of interest to Probe Hybridization Buffer for each embryo sample (final concentration, 10 nM), adding buffer such that the final volume was 200 µL. The samples were moved into the probe solution and incubated for 36-48 hours at 37°C to ensure that the probes fully penetrated the large *He* embryos. The probe solution was removed by washing 4 x 15 minutes in Probe Wash Buffer (Molecular Instruments) at 37°C. Samples were washed 2 x 5 minutes in 5x SSCT (5x SSC and 0.1% Tween 20).

Samples were then pre-incubated in Amplification Buffer (Molecular Instruments) for 30 minutes at room temperature. During this incubation step, DNA hairpins with Alexa-488, Alexa-546, or Alexa-647 fluorophores (Molecular Instruments) and B1, B2, or B3 binding regions were heated to 95°C for 90 seconds and then stored in the dark at room temperature for 30 minutes.

Hairpin solutions were prepared by adding 4 µL of hairpin h1 and 4 µL of hairpin h2 that match the B1, B2, or B3 probe attachment to Amplification Buffer for each embryo sample (final concentration, 60 nmol), adding buffer such that the final volume was 200 µL. Samples were moved into the hairpin solution and incubated for 16-24 hours in the dark at room temperature. Hairpin solution was then removed by washing 3 x 20 minutes in 5x SSCT.

Samples were incubated for 3-6 hours in 1:500 DAPI in 500 mM NaCl solution (500 mM NaCl in 1x PBS) in the dark at room temperature. The embryos were then cleared in glycerol solution by incubating for 1 hour in 50% glycerol in 1x PBS, and then moving to 70% glycerol in 1x PBS. Samples were mounted for imaging on slides in the 70% glycerol solution.

To verify probe specificity, the HCR protocol was run with hairpins only (no gene-specific probes), as well as with no probes and no hairpins. No localized fluorescent patterns were observed under either condition (Fig. S6).

#### Microscopy and image analysis

Mounted embryos and larvae were visualized using a Zeiss 880 Airyscan inverted confocal microscope using a 10x/0.30 EC Plan-Neofluar objective. Images were analyzed using the FIJI distribution of ImageJ (Schindelin et al., 2012). Only linear adjustments, including the “minimum” and “brightness” settings for each color channel, were made to remove background fluorescence. All images presented in this article are single slices of the original multi-slice z-stacks.

## Data Availability

All code used to conduct the analyses and generate the figures in this paper can be found at this repository: https://github.com/brennandmcdonald/He_Larval_scRNA-seq.

## Acknowledgements

We thank Esther Miranda, Zach Pracher, Hannah Devens, Jane Swart, Carl Manner, Emma Wallace, and other members of the Wray lab for their helpful advice during this project. We also thank the staff of the Sydney Institute for Marine Sciences for supporting our data collection efforts in Australia. We would like to acknowledge the use of the Duke University Light Microscopy core facility for preparing micrographs of the HCR samples presented in this paper, as well as the Duke University Sequencing and Genomic Technologies core facility for sequencing the single cell libraries. This work was funded by National Science Foundation grant IOS 1929934 to GAW. BDM also received funding support from the Duke University Undergraduate Research Support Office.

## Supplemental Figure and Table Legends

**Figure S1: Quality control plots for the *He* scRNA-seq atlas.** (A) Violin plot showing the distribution of the number of RNA features (unique gene transcripts) in each cell for each sample time point. (B) Violin plot showing the distribution of the number of RNA transcripts in each cell for each sample time point.

**Figure S2: Dot plot of cell type marker gene expression patterns.** ∼3-6 enriched marker genes (see Table S1) for each cell type in the scRNA-seq atlas were selected, and the average expression of the genes in each cell type was plotted.

**Figure S3: Phylogeny of cell types in the scRNA-seq atlas.** This plot was generated using Seurat’s BuildClusterTree function, which generates the phylogeny using pairwise distances between the cell types in gene expression space.

**Figure S4: Dot plot of neural cell type marker gene expression patterns.** ∼3-6 enriched marker genes (see Table S2) for each neural cell type in the scRNA-seq atlas were selected, and the average expression of the genes in each cell type was plotted.

**Figure S5: Expression patterns of the candidate transcription factors for adult rudiment development.** Each UMAP shows the scRNA-seq expression pattern of one of the 33 candidate transcription factors that was shown to be enriched in adult rudiment tissues (rudiment ectoderm and posterior ectoderm).

**Figure S6: Control experiments to verify HCR specificity.** (A) HCR micrographs of 53 hpf *He* larvae that were incubated with no probes and no hairpins. (B) HCR micrographs of 53 hpf *He* larvae that were incubated with no probes. Neither condition resulted in localized fluorescent patterns, confirming the specificity of the probes used in this study.

**Table S1: Cell type marker genes and references for the full scRNA-seq atlas**

**Table S2: Neural cell type marker genes and references for the neural-only scRNA-seq atlas**

**Table S3: List of putative transcriptional regulators of adult rudiment development Table S4: Probe sequences used in HCR experiments**

